# Mapping Gene-Microbe Interactions: Insights from Functional Genomics Co-culture Experiments between *Saccharomyces cerevisiae* and *Pseudomonas* spp

**DOI:** 10.1101/2020.06.01.127472

**Authors:** Guillaume Quang N’guyen, Mani Jain, Christian R Landry, Marie Filteau

## Abstract

Microbial interactions contribute to shape ecosystems and their functions. The interplay between microorganisms also shapes the evolutionary trajectory of each species, by imposing metabolic and physiological selective pressures. The mechanisms underlying these interactions are thus of interest to improve our understanding of microbial evolution at the genetic level. Here we applied a functional genomics approach in the model yeast *Saccharomyces cerevisiae* to identify the fitness determinants of naïve biotic interactions. We used a barcoded prototroph yeast deletion collection to perform pooled fitness competitions in co-culture with seven *Pseudomonas* spp natural isolates. We found that co-culture had a positive impact on fitness profiles, as in general the deleterious effects of loss of function in our nutrient-poor media were mitigated. In total, 643 genes showed a fitness difference in co-culture, most of which can be explained by a media diversification procured by bacterial metabolism. However, a large fraction (36%) of gene-microbe interactions could not be recaptured in cell-free supernatant experiments, showcasing that feedback mechanisms or physical contacts modulate these interactions. Also, the gene list of some co-cultures was enriched with homologs in other eukaryote species, suggesting a variable degree of specificity underlying the mechanisms of biotic interactions and that these interactions could also exist in other organisms. Our results illustrate how microbial interactions can contribute to shape the interplay between genomes and species interactions, and that *S. cerevisiae* is a powerful model to study the impact of biotic interactions.

## Introduction

The microbiome era has brought upon the importance of prokaryote-eukaryote interactions. Indeed, we have come to realize that animal and plant-associated microbiomes contribute to the functions and health of their host, and thus to their evolution (1-3). On the microbial scale, the subclass of bacterial-fungal interactions has also been deemed crucial to the functions of many ecosystems, contributing to biogeochemical cycles, biotechnology, food production, as well as plant, animal and human health and development (4). Thus, interactions between prokaryotes and eukaryotes are fundamental to most ecosystem, yet much remains to be uncovered about the molecular basis of interkingdom signaling (5).

Bacterial-fungal interactions could make excellent models to study prokaryote-eukaryote interactions (4). Indeed, bacterial-fungal associations could be used to assess the evolutionarily conserved molecular mechanisms between prokaryote and eukaryotic cells (4). A variety of physical and molecular interactions have been reported between fungi and bacteria resulting in different outcomes for each partner (6). High-throughput methods have been developed to identify metabolites involved in these interactions (4), but less attention has been given to the genomic elements involved. Moreover, little is known about the fungal counterpart of these interactions at the mechanistic level (7).

The model yeast *Saccharomyces cerevisiae* would be an ideal system to study the genes and functions that underly bacterial-fungal interactions. Owing to multiple concerted efforts to systematically screen deletion mutant phenotypes (8) and genetic interactions (9), a lot is known about the metabolic and cellular function of the budding yeast genes. Yet over 700 of the ORFs are still uncharacterized in terms of biological function (10). The cellular mechanisms underlying the strategies used by *S. cerevisiae* to interact with other microorganisms in various conditions remain largely unknown (11). Only three yeast genes (MAK32, MKT1 and ATF1) are annotated with the ontology term “interspecies interaction between organisms” (10). Systematic surveys of gene-microbe interactions are lacking and could provide new insight into these ORFs functions. Moreover, a pangenome analysis of *S. cerevisiae* reports a strong functional enrichment for cell-cell interactions for ORFs whose presence is variable in the population (12).

The yeast genetic targets of some antimicrobial compounds produced by other organisms has been studied using chemogenomic approaches including barcode sequencing (Bar-seq) (13, 14). However, these approaches consider only one-sided effects of single molecules. Instead, co-culture experiment may not only demonstrate interactions that are the result of specific antimicrobial compound activity, but also indirect interactions that are metabolic in nature or that are the result of cell-to-cell contact. Indeed, some *S. cerevisiae* strains have been shown to interact with bacteria both indirectly by the production of antimicrobial peptides (AMPs), as well as by metabolic cross-feeding and directly by physical contact (11, 15). Most of the interspecies interaction knowledge at the genomic level in *S. cerevisiae* comes from transcriptomic co-culture experiments (16-19), which does not allow to measure the impact on fitness as it captures only the expression level at the moment of sampling. Combining bar-seq and co-culture experiments would thus allow to measure the impact of complex biotic interactions and identify the fitness determinant involved.

To capture ecologically relevant interactions, natural models that simulate plausible encounters are needed. Moreover, novel interactions make a relevant model system as they can occur when cells meet in the context of an ecological succession where new niches are formed such as in foods, fresh wounds, or newborn hosts (20). *Saccharomyces* yeast and *Pseudomonas* bacteria co-occur in several environments and can form biofilms e.g., in kefir (21), olives (22), tree bark (23), wine fermentation (24) and winery wastewater (25) and thus, make an ecologically relevant pairing. Here we coupled co-culture and bar-seq experiments to study microbial interactions between the model yeast *S. cerevisiae* and seven *Pseudomonas spp*. isolated from maple sap. Since *S. cerevisiae* is not naturally found in maple sap (26), this pairing constitutes a novel interaction that may reflect the diversity of mechanisms at play between new partners as well as evolutionarily conserved pathways involved in interkingdom signaling.

## Results

Seven *Pseudomonas spp*. strains isolated from maple sap, with different negative impact on fungal growth on solid media against *S. cerevisiae* were used to investigate gene-microbe interactions (Table 1). Sequences of the 16S rRNA and the RpoB genes classify them in the *Pseudomonas fluorescens* complex of species.

**Table 1.**
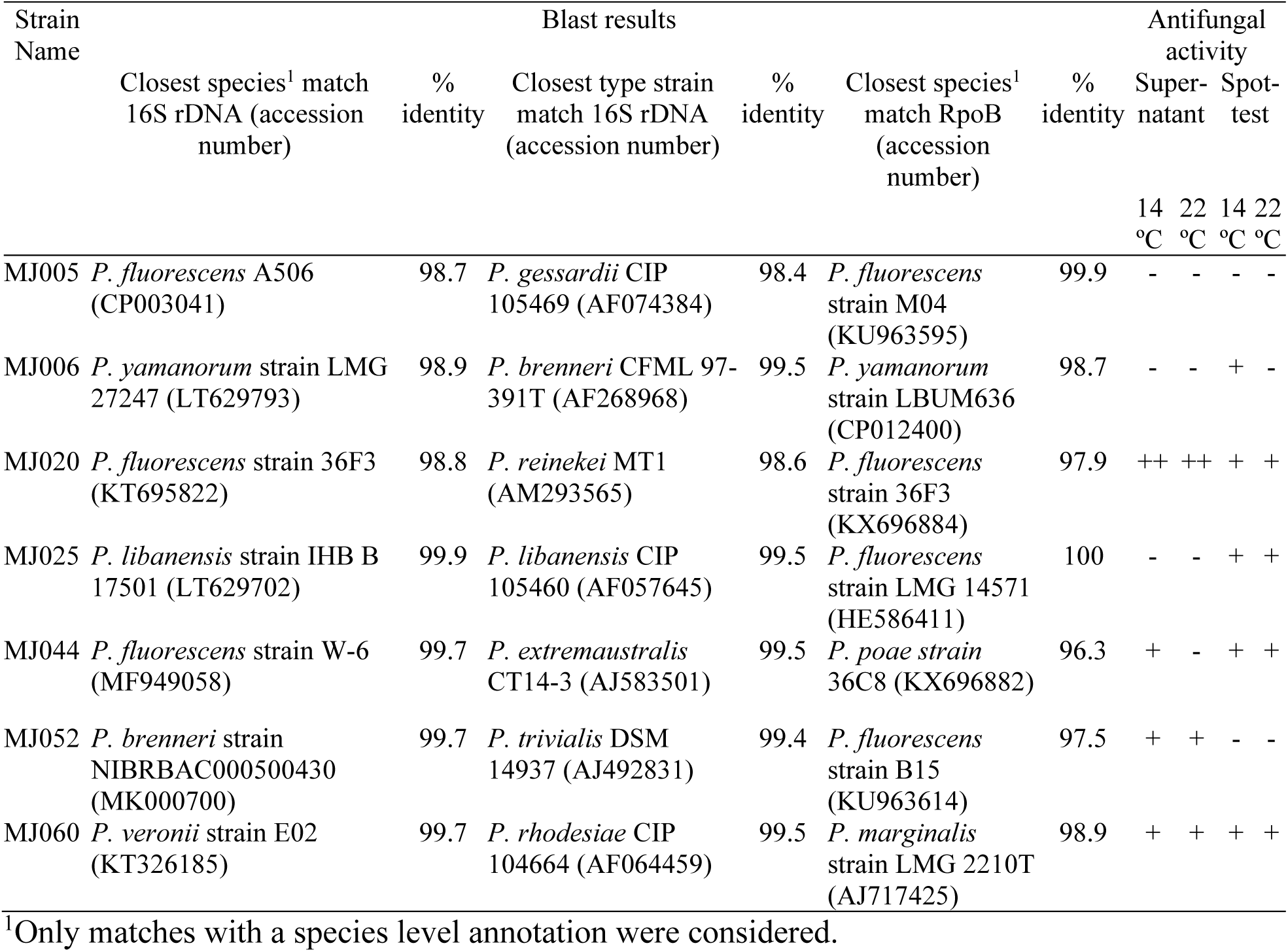
Phylogenetic affiliation of the *Pseudomonas* strains.

We used these *Pseudomonas spp*. strains to perform a functional genomic co-culture experiment with a prototrophic version of the *S. cerevisiae* deletion collection. The co-cultures were maintained for four 24h growth cycles in a synthetic allantoin media (SALN) simulating maple sap along with a control condition consisting of the *S. cerevisiae* deletion collection alone (Fig. 1a). Yeast growth at the end of each cycle remained around 10^7^ cell/mL. Co-cultures with MJ005, MJ006, MJ020 and MJ052 strains showed lower counts than the control after each cycle. MJ025 and MJ044 also had lower counts than the control at cycles 1 and 3. This difference indicates that some level of competition or inhibition of yeast growth took place during the experiment. The MJ060 co-culture did not follow the same trend, having higher counts after the first cycle, then lower counts in subsequent cycles. Thus, this co-culture exhibited globally both positive and negative interactions with the yeast deletion pool. After the fourth growth cycle, the fitness of each yeast deletion strain was measured by barcode sequencing.

**Figure 1.**
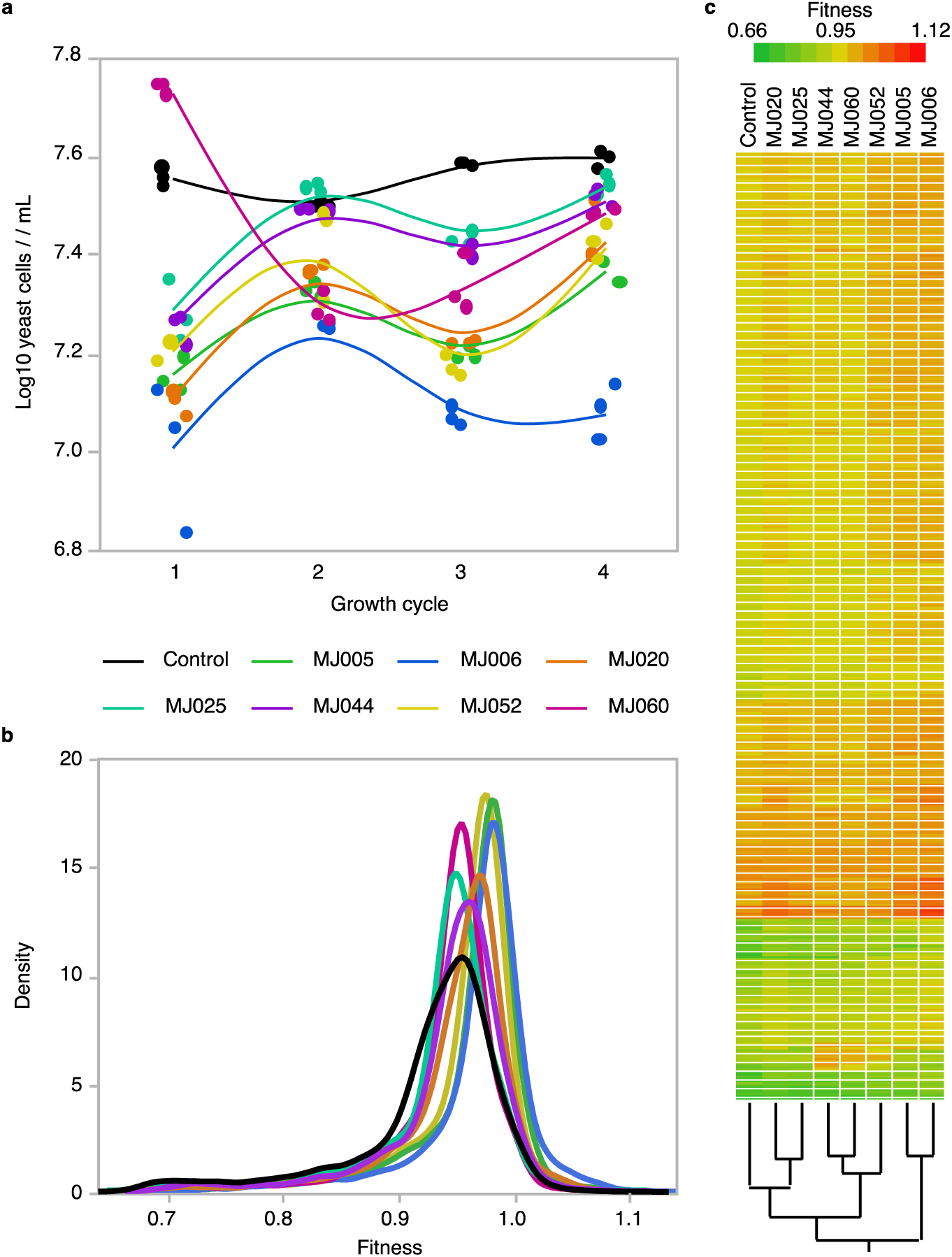
Co-culture experiment between seven *Pseudomonas* spp. strains and the pooled *S. cerevisiae* strain deletion collection. a) Counts of yeast cells after each cycle of 24h co-culture. b) Density curve of the yeast deletion strain fitness distribution for each co-culture compared to the *S. cerevisiae* strain deletion collection alone (Control). c) Hierarchical clustering and heatmap of averaged fitness profiles of control and co-cultures.

We estimated the absolute fitness of 4 285 deletion strains relative to the average of strains with a pseudo-gene deletion as an internal wild type control in the pool (See methods, supplementary file 1). All fitness distributions in co-culture were significantly different from the pure culture control (Kolmogorov-Smirnov Asymptotic Test, Prob > D+ <0.0001) (Fig1b). The dispersion of fitness as estimated by the variance was narrower for all co-cultures than for the control (Dunnett’s test, P-value<0.01) indicating that the fitness was more homogenous among the deletion strains. The co-cultures had a higher average fitness (ranging between 0.94 and 0.97) compared to the control (0.93), but the difference was only significant for MJ005, MJ006, MJ020 and MJ052 (Dunnett’s test, P-value <0.05), meaning that in general the deleterious effects of loss of function were mitigated in co-culture. The average fitness of the yeast deletion pool negatively correlated with overall yeast growth in our experiments (Spearman P = -0.8548, P-value = 1.6e-28) reflecting that stronger selective pressures on the yeast pool, either by competition or inhibition exerted by the bacteria, yielded an overall fitter yeast population. When comparing the fitness profiles of each co-culture and the control by hierarchical clustering, the grouping did not follow the average fitness pattern, indicating that different mechanisms of interaction are also at play at the genetic level (Fig. 1c).

To characterize the impact of co-culture on deletion strain fitness while accounting for whole population shifts, we used the residual difference of a linear fit to the control. We then performed a functional enrichment analysis on the ranked residual fitness (Fig. 2, supplementary file 2). We found significant (FDR<0.01) enrichments for fitness loss and gain in each co-culture compared to the control. A lower fitness in co-culture means that the deleted gene contributes to a function under selective pressures, thus contributes to competition or avoidance of inhibition. Accordingly, the top fitness decrease enrichment in most co-culture was related to specific resources, either oxygen (MJ044, response to decreased oxygen levels), nitrogen (MJ020, MJ052 and MJ060, allantoin catabolic process), carbon (MJ005, maltose metabolic process) or vitamin (MJ006, nicotinamide nucleotide biosynthetic process), suggesting competition. The main carbon source in our media was sucrose, a disaccharide that can also be the substrate of enzymes in the maltose metabolic pathway. Conversely, a higher fitness in co-culture may be the result of relaxed selective pressures, allowing the strains to recover fitness if the deleted genes contribute to a function that is no longer necessary, or if their deletion allows to evade inhibitory mechanisms. Fitness gains processes were enriched for mitochondrial gene expression (all co-cultures but MJ044) (Fig. 2). Among the strains associated with mitochondrial gene expression in our data set (Supplementary file 2), a few strains (14/83) actually improved their fitness relative to the wild type (>1) in co-culture (Supplementary file 1), but the majority only partly recovered from the deleterious effect of gene loss (Fitness <1), suggesting a general relaxation of the importance of mitochondrial functions in co-culture. However, we also find a fitness gain enrichment for tRNA wobble uridine modification in some co-cultures (MJ005, MJ020 and MJ060) that could represent the evasion of inhibitory mechanisms, since the deletion of genes in this process is known to confer resistance to microbial toxins (27).

**Figure 2.**
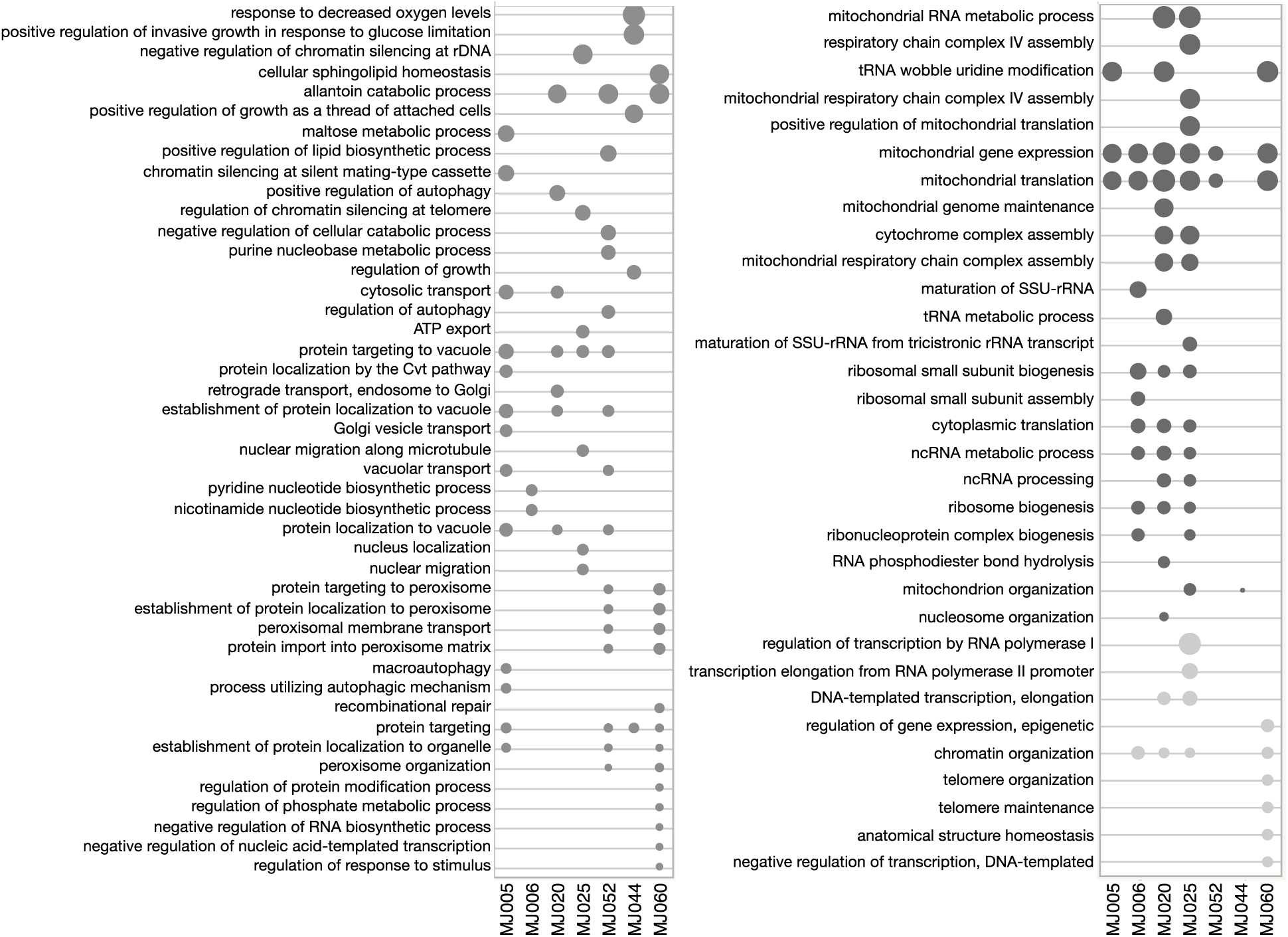
Gene Ontology Process enrichments of ranked fitness residuals in co-culture experiments. Enrichments were performed with the online tool of the String Protein-Protein Interaction Networks database (https://string-db.org). The left panel shows enrichment for the top of the list corresponding to a fitness decrease in co-culture. The right panel shows the enriched processes of the bottom of the list corresponding to a fitness increase in co-culture (dark grey dots) and those enriched from both ends (light grey dots). Dot size is proportional to the enrichment score (FDR <0.01). See Supplementary file 2 for complete results.

An increase in fitness in co-culture compared to the control could be attributed to the indirect effect of the metabolic impact of the bacteria on its environment, resulting in cross-feeding wherethe bacteria release metabolites that complexify the nutrient-poor media used in our experiments. Thus, we would expect an overlap with strains that are affected by media composition, for instance, those that show a fitness difference between a defined synthetic media and a complex rich media containing yeast extract such as YPD. To validate this assumption, we first defined a list of deletion strains for which there was a significant fitness difference (Dunn test, FDR adjusted P-value<0.05) between a co-culture and the control and a minimum residual difference of >|0.02| of the linear fit to the control. In total, 643 yeast deletion strains were identified, including strains that were specific to each *Pseudomonas* co-culture (Fig. 3a). In fact, most deletion strains were identified in a single co-culture (364 out of 643 in total). We then compared these lists with the results obtained by (28) in synthetic complete (SC) and complex media, either based on a fermentable carbon sources (YPD) or requiring respiration (YPG). We found that indeed our candidates are enriched for strains whose fitness is different between SC and YPD (Fisher’s exact right test, P-value= 1.6e-18), but also between YPD and YPG (Fisher’s exact right test, P-value= 2.6e-29). Taken together with the functional enrichments highlighted above, this result underscore the importance of mitochondrial functions in our bacterial-fungal co-cultures. Moreover, the overlapping strains tend to be identified in multiple co-cultures (Likelihood ratio P-value 1.6e29) indicating that enrichment of the media may be a common occurrence in co-culture (Fig. 3b).

**Figure 3.**
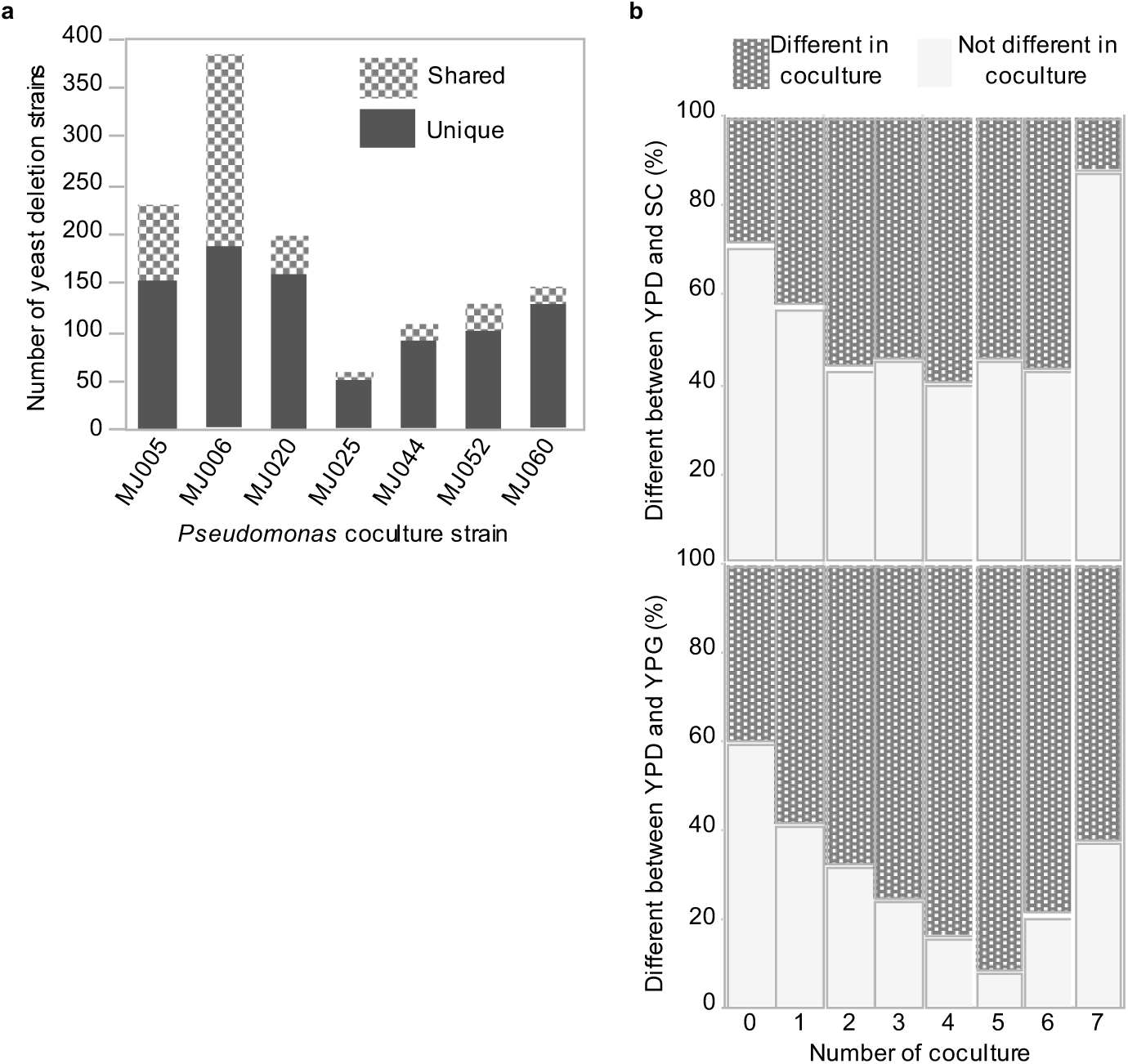
Yeast deletion strains showing a significant difference in fitness when co-cultured with a *Pseudomonas spp*. strain. a) Number of yeast strain identified in each co-culture. b) Percent overlap between strains identified in our co-cultures and strains showing a fitness difference between a synthetic culture media (SC) a fermentable complex media (YPD) and a non-fermentable complex media (YPG), classified by the number of co-cultures in which the strains were identified.

To further ascertain the extent to which the effects observed in co-culture were the result of direct or indirect interactions, we retrieved the yeast fitness profiles in various maple sap samples from a previous experiments using similar methods (29). Correlations between these maple sap fitness profiles and co-culture results in our control media, which simulates sap, are strain dependent (Fig 4). MJ052, MJ044 and MJ060 show significantly higher correlations (Dunnett’s Method, P-value <0.0001) to maple saps than the control profile, while MJ006 and MJ020 correlations are significantly lower (Dunnett’s Method, P-value <0.0001). Thus, the effect of some co-culture on synthetic sap can mimic to some extent the effects observed on natural maple sap. This further supports the idea that the main impact of these strains on yeast fitness is indirect and metabolic in nature. For other co-cultures however, the effects on fitness profiles across genes appear to diverge from maple saps, indicating that other mechanisms may be at play, such as cell-cell interactions, or that these isolates do not contribute to maple sap properties.

**Figure 4.**
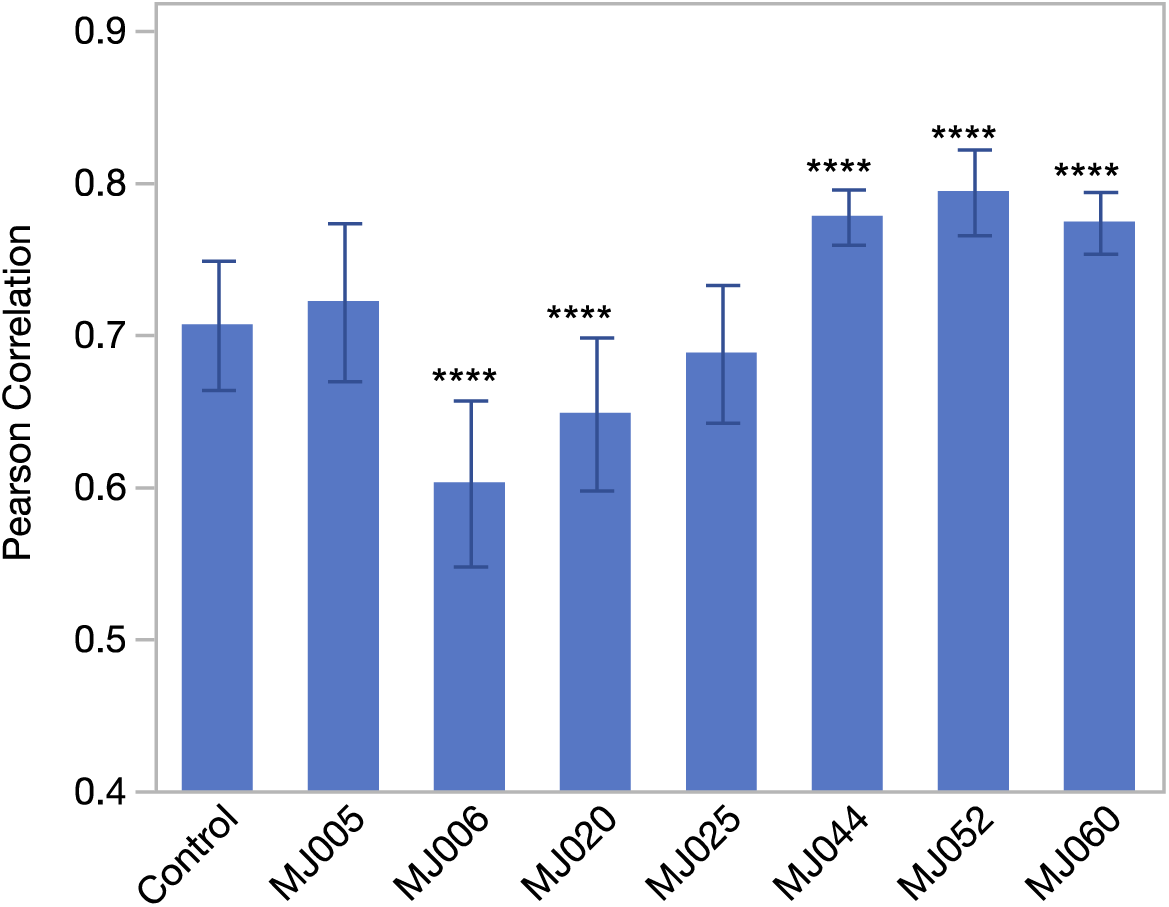
Correlation between yeast fitness profiles in co-culture with *Pseudomonas* spp strains and yeast fitness profiles in 27 samples of maple sap. Bars represent the average Pearson Correlation and error bars the standard deviation. **** indicate a significant difference with the control (Dunnett’s test, P-value<0.0001).

To assess if fitness effects were specifically triggered in co-culture, we measured the growth of 11 yeast deletion mutants in *Pseudomonas* spp. cell-free supernatants. Out of the 77 combinations tested, 49 (64%) where coherent in their fitness effect sign (Fig. 5). Sign inconsistencies were observed at least once for each deletion mutant and each *Pseudomonas spp*, however the *drs2* mutant gave the most divergent results, along with MJ005 and MJ060. Notably, the *drs2* mutant had a negative fitness in all co-cultures, but a fitness gain in all but one supernatant, indicating that Drs2 could play a positive role in the direct or feedback interactions with *Pseudomonas*.

**Figure 5.**
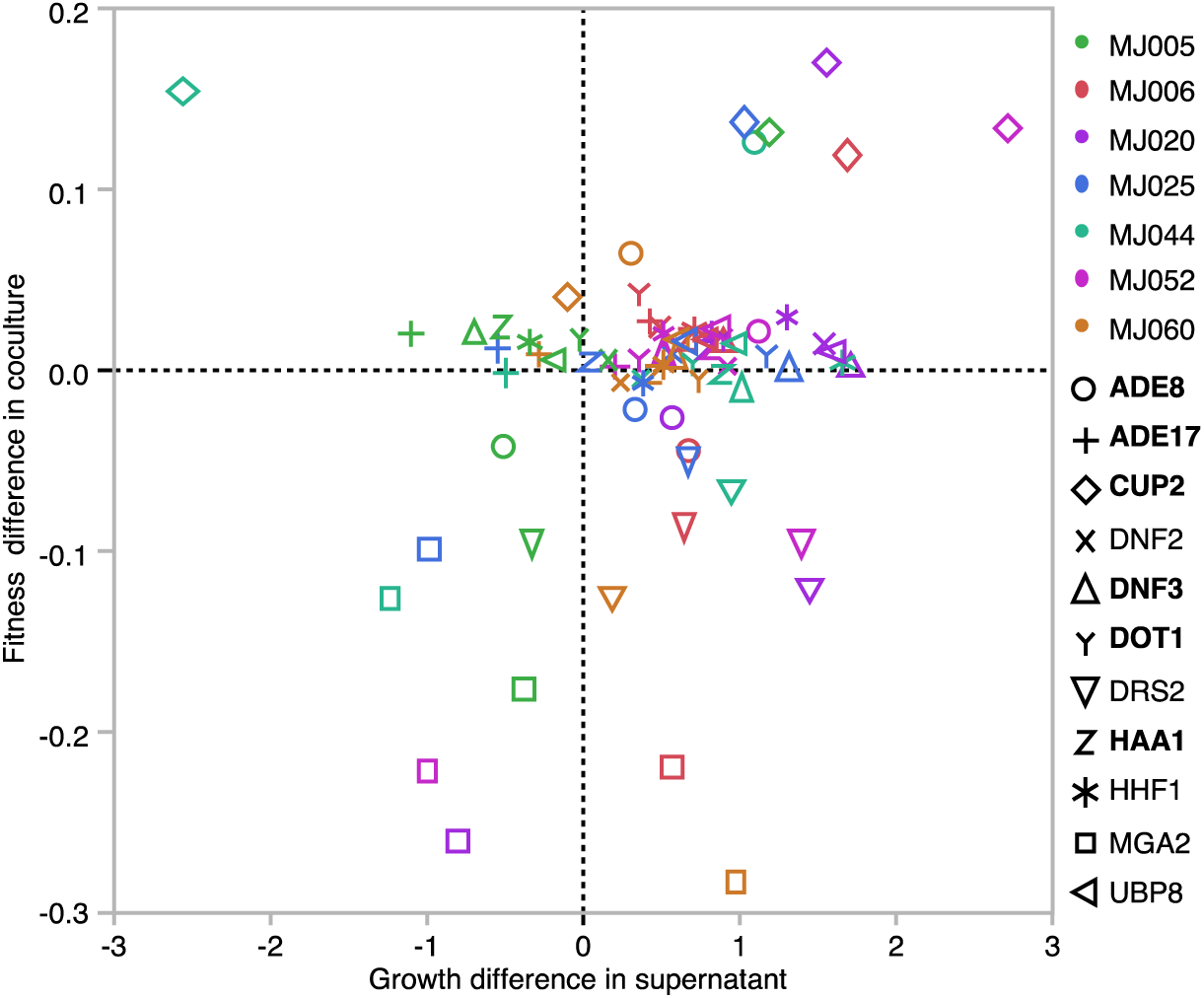
Relative impact of *Pseudomonas spp*. co-culture compared to supernatant exposition on yeast deletions mutants. *Pseudomonas* strains are identified with colors, and yeast deletion strains by symbols.

We hypothesized that some yet uncharacterized ORFs may be specifically involved in interspecies microbial interactions and thus those would be revealed in our experiment. We found that uncharacterized ORFs are underrepresented in our co-culture lists compared to verified ORFs (Fisher’s exact test right P-value=1.12e-7). Nevertheless, 43 strains deleted for an uncharacterized ORFs show fitness differences in our co-culture experiments and nearly half of those (19/43) have also been reported to be differentially expressed in *S. cerevisiae* when co-cultured with either *Torulaspora delbrueckii, Candida sake, Hanseniaspora uvarum, Saccharomyces kudriavzevii or Oenococcus oeni* (Figure 6). Given that the phenotypes assayed, the culture media and the co-culture microorganisms were vastly different, the overlap of these findings suggest that these uncharacterised genes may play a generic role in mediating interspecies interactions.

**Figure 6.**
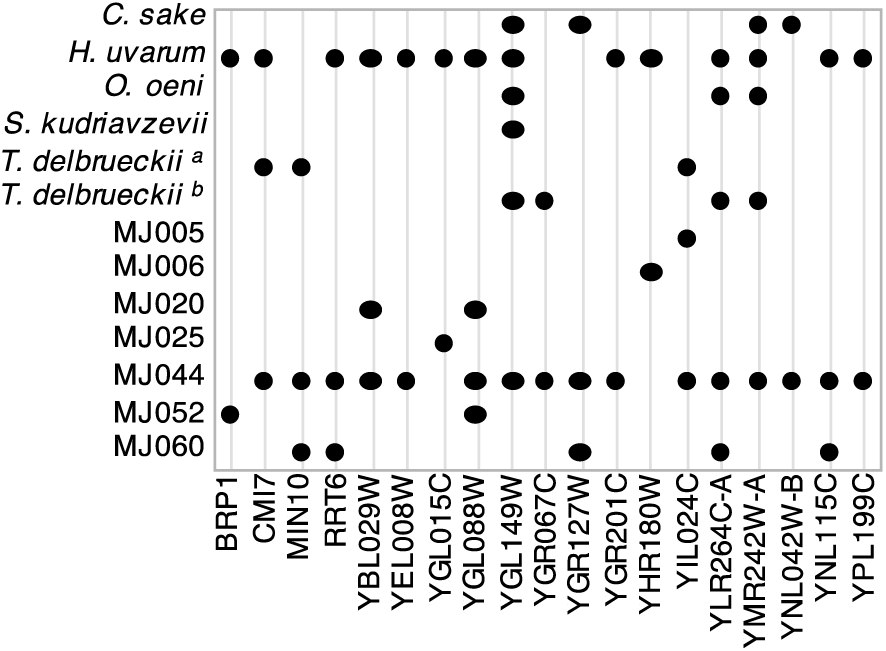
Unknown function genes identified in our co-culture experiments and other interspecies co-culture studies with *C. sake* (17), *H. uvarum* (17), *O. oeni* (18), *S. kudriavzevii* (16) *or T. delbrueckii* ^a:^(17)^, b:^(19).

To assess the phylogenetic specificity of our gene-microbe interactions, we looked for enrichments for homologs in other eukaryote species in our co-culture gene lists (Fig. 7). We found that six co-cultures show such enrichments. Also, genes with homologs in the *Candida* genus appear more frequently than expected by chance in most co-cultures (MJ006, MJ020, MJ025, MJ052 and MJ060). Interestingly, MJ006 and MJ020 gene-microbe interactions are enriched for genes conserved in both fungi and metazoan. Functional enrichment analysis of REACTOME pathways using ranked fitness residuals reveal that deletion strains have a fitness gain in these two co-cultures in pathways conserved in humans such as eukaryotic translation initiation (Supplementary file 2). Thus, the mechanisms involved in these bacterial-fungal associations may extend to other bacteria-eukaryote pairings.

**Figure 7.**
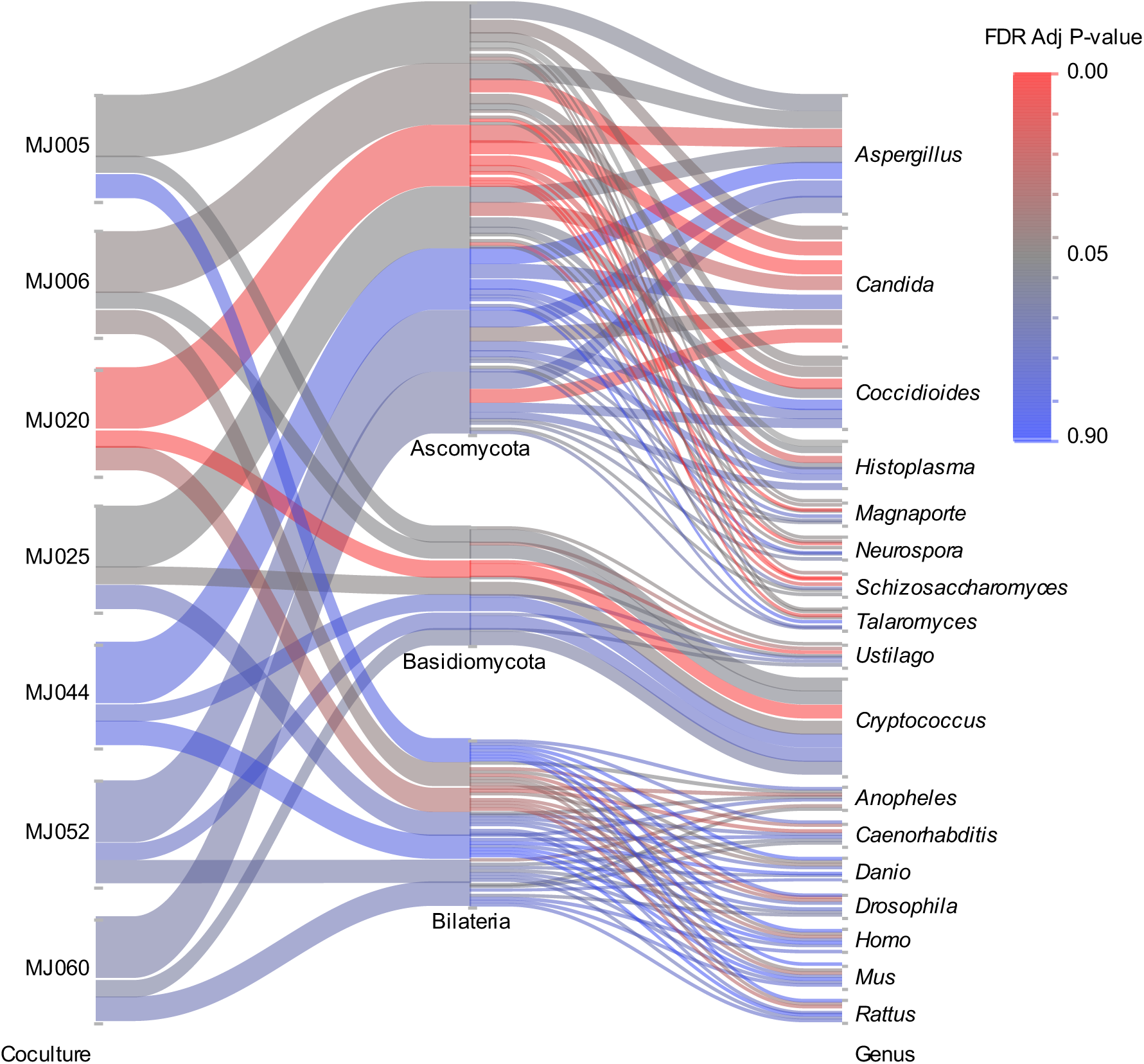
Enrichments for homologs in other eukaryotes in our co-cultures gene lists. Link colors show enrichments by taxa for each co-culture. Line thickness reflects the number of reference organisms within each level used to calculate the average enrichment.

## Discussion

Interspecies interactions are an important ecological process, yet the qualitative and quantitative impact of biotic interactions at the genetic level i.e., gene-species interactions remain largely unknown. To characterize these effects in a model system, we studied a naïve interaction, where partners have not met before, between *S. cerevisiae* and seven *Pseudomonas spp*. strains. In functional genomic screen co-cultures, we observed mostly negative effects on overall yeast growth. Different types of interactions can underlie these effects such as competition or inhibition (amensalism). Based on the preliminary antifungal assays on solid media, yeast inhibition was expected in most cases. Competition for resources was also expected because they are common in carbon-rich environments (30). However, the positive effect of MJ060 after the first day of the experiment was unexpected. Either mutualistic, synergistic or commensalistic relationships could explain this result, but require further investigation to be distinguished. Because our data reflect the net effect on fitness after several transfers and generations, this effect may be diluted in our results.

Co-culture of *S. cerevisiae* with *Pseudomonas* spp. induced a decrease in fitness diversity and an increase in average fitness which is indicative of the combined biotic selection forces at play. In yeast, it was previously reported that gene loss is in general slightly deleterious in laboratory conditions (31). Here, the co-culture with bacteria partly rescues the deleterious effects of gene loss. We also observed that stronger competition or inhibition effects result in a higher average fitness of the yeast population. Moreover, overlap with results from another study comparing synthetic to complex media (28) suggest a metabolic effect via media complexification by the bacteria. Taken together, these results point to nutrient cross-feeding interactions, that is when one microorganism provides a nutrient to a partner, thus relaxing its metabolic burden (32). Moreover, a significant overlap with results diverging between a fermentable (glucose) to a non-fermentable (glycerol) substrate was observed. In fact, strains harboring deletion of genes encoding mitochondrial functions appear to recover fitness in co-culture with the *Pseudomonas* strains. This can be explained if these functions are no longer advantageous given the conditions available. For instance, in addition to respiration, mitochondrial function contribute to heme synthesis and amino acid metabolism in yeast (33). Meanwhile, the presence of exogenous co-factors such as heme can influence respiratory metabolism (32). Also, cross-feeding can noticeably be predicted in computationally designed media where amino acids are absent (34), such as in our experiment.

Alternatively, a fitness gain in co-culture could be explained if mitochondrial functions are the target of an antimicrobial mechanism. For example, the *Pseudomonas* strains could produce compounds that block oxidative respiration in the wild type, thus allowing deficient mutants to recover fitness in the population. This particular mode of action has been observed for 4-hydroxy-2-heptylquinoline N-oxide (HQNO) and pyocyanin produced by *P. aeruginosa* (35). We also identified gene involved in tRNA wobble uridine modification as fitness determinants, which are known targets of microbial toxins (27).

It is noteworthy that the effect on the overall yeast growth over the experiment is similar between several co-cultures showing competition or inhibition, yet the resulting fitness profiles are quite different. While it is expected that microbial interactions are condition-dependent (34), our results show that the outcome is also species-dependent, even among closely related bacteria. In fact, we find evidence of competition for different resources, either oxygen, nitrogen, carbon or vitamins, depending on the *Pseudomonas* strain. Hence, even though the culture media is the same, the bacteria do not impact the same genomic fitness determinants in yeast, meaning that these players do not necessarily adopt the same game strategies. A similar conclusion was reached with lactic acid bacteria that adopted different strategies in the face of interspecies competition (36).

Overall, the fitness patterns of loss and gain in our co-cultures imply some balance between competition and cross-feeding. The MJ044 co-culture appears as the exception, where we find evidence of competition for oxygen i.e., deletion of genes involved in the response to decreased oxygen levels lead to decreased fitness in co-culture, but no fitness recovery for mutants associated with mitochondrial gene expression. Altogether, these observations underscore the importance of biotic selective pressures over environmental conditions. In a complex community, cells would then be challenged by multiple selective pressures at the same time which could highly constrain the evolutionary trajectory.

The fitness determinants in *S. cerevisiae* when grown in maple sap as a natural substrate have previously been identified (37). The co-cultures were performed in an allantoin and sucrose based synthetic culture media that has been shown to mimic maple sap properties in term of growth parameters of a wild *Saccharomyces paradoxus* population (37). Our result show that some *Pseudomonas* strain when co-cultured in this synthetic sap media can recapitulate to some degree the effect on yeast fitness observed in natural maple sap samples. Commercial maple sap has been used as a natural source of nutrient to study mold metabolism and development (38). However, maple sap is unavoidably contaminated, mostly by bacteria and yeast during its collection through complex networks of plastic tubing (26) and then sterilized before it can be sold or used as a culture media. Therefore, maple sap is a natural vegetal substrate, but also to some degree a spent (or fermented) culture media, which can explain in part its variation in composition. Thus, our results illustrates the important contribution of biotic effects in natural environments, even if they are indirect.

Allantoin is an ureide that can be used as a sole source of nitrogen by *S. cerevisiae* and peculiarly, genes required for the degradation of allantoin (DAL) form one of the two metabolic genes clusters found in its genome (39). Our experiments revealed that MGA2, which is localized next to the DAL cluster contributes to yeast fitness in co-culture with all *Pseudomonas spp*. tested and that this effect is mediated via a bacterial metabolite in at least five cases since exposition to the supernatant of the bacteria is sufficient to reproduce the fitness defect in the mutant. The molecular function of MGA2 is still unknown, but it is involved in transcription regulation in response to hypoxia and iron starvation (10).

In contrast to MGA2, when testing for the effect of supernatant only, results for the DRS2 mutant showed the most sign-inconsistency compared to the co-culture fitness competition. The lack of supernatant effect indicates that the particular mechanism underlying the fitness effect on this mutant is not mediated by a constitutively secreted metabolite. Our co-culture approach has the potential to capture other mechanisms of action as well, involving an induction/feedback mechanism, a physical contact, or even a higher-order interaction with other yeast in the deletion pool. Indeed, DRS2 is annotated with response to pheromone triggering conjugation with cellular fusion (10). Further experiments would be needed to discriminate between these possibilities, such as co-culture in membrane-separated wells (40). Also, further interpretation of the effects observed would be possible by incorporating genomic and transcriptomic information about the bacterial competitors. For instance, most members of the *P. fluorescens* and *P. gessardii* phylogroups possess the *HicA* gene, which encodes a toxin acting by inhibiting translation, while it is absent from the *P. jessenni* and *P. koreensis* groups (41).

Our approach highlights genes for which a loss of function is compensated or aggravated in co-cultures with bacteria. Some of these gene-microbe interactions may be specific to the model microorganisms used, but others may extend to additional organisms where orthologs of the target genes are present. On the one hand, some genes may play a role in unspecific interspecies interactions like competition sensing. Indeed, we find interspecies interaction for 19 unknown function genes in common with other studies (16-19). On the other hand, some interactions may be specific to the organisms involved and depend on their life history. The r/k-selection is an ecological theory that could help explain the different pattern of co-culture effects observed. In this framework, r-strategist microorganisms are fast growing generalist thriving when nutrient are plentiful, while k-strategist are competitors specialized for specific or limited resources (20). In a similar interpretation, r-strategist would engage in generalist interactions, targeting broadly conserved functions or genes, while k-strategist would have more specific interactions mechanisms. Surprisingly, all except the MJ004 co-culture revealed gene-microbe interactions that involve genes conserved in other eukaryotes. Genes conserved in the *Candida* genus were enriched in these six co-cultures. The *Pseudomonas* strains that were used were all isolated from maple sap were several groups of yeast can be found, including *Candida sake* (26). However, the *Pseudomonas* strains may have been introduced in this environment at different times and may have different life history traits that shaped their behavior in co-culture. For instance, we find that MJ006 and MJ020 affects genes that are preserved among eukaryotes and enriched for eukaryotic translation initiation. The significance of this result about how it relates to the life history of the strain can only be hypothesized at this point. Because co-culture with these two *Pseudomonas spp*. do not recapitulate growth in maple sap, it may be hypothesized that they were introduced in the sap during collection by insects or human manipulations, but are not endogenous to the maple phyllosphere, or that they simply are r-strategists. The various patterns of enrichment for conserved functions reported here highlight that our approach could be used to answer questions about the relationship between life history and the evolution of interactions mechanisms. It would also be interesting to test if some gene-microbe interactions identified here are conserved in higher eukaryotes. In this context our approach could be useful to study host-microbiota interactions on a genomic level.

The molecular mechanism of a specific positive interaction between *S. cerevisiae* and *Pseudomonas putida* in grape juice was revealed using a phenotypic screen with the yeast deletion collection (7). The study identified the mode of action of a phenotypic change in *P. putida* induced by the yeast, demonstrating the usefulness of the deletion collection to study specific microbial interaction mechanisms. Here, we used the yeast deletion collection in a liquid competition assay to reveal the fitness determinant involved in naïve interactions without a priori knowledge of their impact. Thus, our approach is generalizable to other pairings. Given the plethora of tools developed for *S. cerevisiae*, such as mutant collections (42-45), including humanized yeast (46), the approach presented could find applications in the study of interactions with microorganisms of clinical or technological importance such pathogens or biocontrol agents.

In conclusion, our study demonstrate that *S. cerevisiae* is an excellent model organism to study biotic interaction at the genomic level and further our understanding of the evolution of these interactions. Indeed, this type of bacterial-fungal interaction could be a useful model system to analyze complex interactions (4) and complement other emerging co-culture approaches on model microbiota (47) and experimental evolution of engineered interactions (48).

## Methods

### *Pseudomonas* spp. strains

*Pseudomonas* spp. strains were isolated from maple sap. Maple sap concentrated by membrane processing were obtained from Quebec producers during the spring of 2016. Samples were kept frozen at -20°C until analysis. The sap concentrates were centrifuged for 5 min x 915 g at 4°C and the supernatant was extracted and sterilized by 0.2 µM filtration. Autoclaved water was added to the pellets and 5 µL of each cell culture was plated using glass beads on sterile maple syrup media (37) with or without chloramphenicol (12,5 μg/mL). After 5 days of incubation at 10°C, colonies were streaked on Synthetic Allantoin (SALN) media (1.75 g.L^-1^ of yeast nitrogen base, 1.25 g.L^-1^ of allantoin, 2% sucrose and 2% agar). Stock cultures were prepared in SALN liquid media (without agar) and with 20% glycerol (V/V), then stored at -80°C. Strains identification was performed by sequencing the 16S rRNA subunit gene with universal primers and the RPO-B gene with LAPS_F (TGGCCGAGAACCAGTTCCGCGT) and LAPS27_R (CGGCTTCGTCCAGCTTGTTCAG) primers (49).

### Antifungal assays on solid media

For the seven *Pseudomonas* strains, two antifungal activity testing methods tests were performed on *S. cerevisiae* (*hoΔ::KanMX*), a supernatant assay and a spot-test assay. First, all strains were pre-cultured overnight (16h) in liquid SALN media at 22°C without agitation. For the supernatant assays, the saturated *Pseudomonas* pre-culture were diluted at 1/1000 v/v in 65°C (SALN) media (0.75% agar). This step killed the bacterial cells, which are sensitive to heat. Then, 5 µL of 10-fold serial dilutions starting at 0.1 OD of *S. cerevisiae* culture were deposited on the solidified media. Controls were done without *Pseudomonas* in the media and growth was visually compared between the treatment and the control after 2 to 4 days of incubation at 14°C and 22°C. For the spot-test assays, 50 µL of 0.5 OD *S. cerevisiae* culture were spread on solid SALN media (2% agar) using sterilized beans. Then, 5µl drops of *Pseudomonas* cultures were deposited on the media with concentrations adjusted at ½ and ¼ of the saturated culture. After 2 to 4 days of incubation at 14°C and 22°, *Pseudomonas* strains presenting an inhibition halo were considered having an antifungal effect.

### Cytometry

Cell counts were obtained by flux cytometry in a Biorad flux cytometry Guava easyCyte 14HT flow cytometer (EMD MILLIPORE Ltd, Oakville, ON, Canada). Counts were obtained from 5,000 events of serial dilutions of the co-cultures in a 96 deep-well plate. Size discrimination was for 10^2^ to 10^3^ nm for bacterial strains and 10^4^ to 10^5^ nm for *S. cerevisiae*.

### Co-culture functional genomic screen

The yeast deletion collection used was obtained from (45). The functional genomic screen was performed as in (37), with the following modifications: liquid assays were carried out in 4 mL of SALN media in 24 deep-well plates at 22°C without agitation. Co-cultures between the *S. cerevisiae* deletion collection and the seven *Pseudomonas spp* strains along with a control of the deletion collection alone were performed in triplicate. The *S. cerevisiae* initial OD were adjusted to 0.05 and the *Pseudomonas / S. cerevisiae* ratio were 1/50 for MJ025, MJ006, MJ060, MJ044 and 1/250 for MJ005, MJ020 and MJ052 strains. These ratios were set according to their growth rate to allow sufficient yeast growth. After 24h incubation, 200 µL of co-culture were transferred to 3.8 mL fresh media, for a total of four days. The cell counts were measured after each 24h growth cycle. After four growth cycles, DNA extractions were performed individually on each well. A total of 24 libraries, were then constructed by PCR using the primers listed in Supplementary file 3. DNA extractions, PCR, library preparations and sequencing were performed as in (29).

## Sequencing analysis

Sequencing results were analyzed as in (37), at the Plateforme d’Analyses Génomiques of the Université Laval (IBIS), with the following modifications: the sequence reads were mapped to the reference using Geneious R9, with the following parameters: minimum overlap 113, word length 20, index word length 15, maximum mismatches per reads 6%, allow gap 4%, maximum gap size 2, and maximum ambiguity 16. We obtained 45 122 440 reads after size filtering and 91% of those were assigned unambiguously. As we used the same initial deletion collection pool as in (29), we used those sequencing results in our analysis. We used the 4 285 strains that had a sum of more than 50 reads in the three initial pool replicates for further analysis. The absolute fitness was obtained as in (28) using the average of six strains deleted for pseudogenes (YIL170W, YCL075W, YIL167W, YIR043C, YIR044C and YCL074W) as the wild-type reference.

### Candidate lists enrichment analysis

Statistical and multivariate analysis was performed in JMP13 (SAS Institute, Cary, NC). Dunn tests comparing the fitness of each strains in co-cultures vs in the control *S. cerevisiae* culture were performed to identify significant changes. Threshold for candidates were set at FDR adjusted P-value<0.05 and a residual fitness of a linear fit to the control >|0.02|. Overrepresentation of Gene Ontology and REACTOME pathways were tested using with the online tool of the String Protein-Protein Interaction Networks database (https://string-db.org). The overrepresentation was calculated using the ranked list of co-culture fitness residuals of a linear fit to the control.

Phenotypes for null mutants in the S288C genetic background retrieved from the Saccharomyces Genome Database (10). Gene homologies between *S. cerevisiae* and other species were retrieved from Yeastmine (50). GOSlim annotations were retrieved from the Saccharomyces Genome Database (10). Fitness results from (28) were obtained for the strains of the auxotroph deletion collection grown in various culture media. Since no replicates were available except for YPD, we simply compared our candidate genes to those that show a fitness difference >|0.02| between conditions. Overrepresentation of candidate genes was tested with Fisher’s exact test and FDR-adjusted P-values are reported.

### Supernatant growth experiments

We selected yeast deletion strains (DRS2, CUP2, MGA2, HHF1, UBP8, DNF2, DNF3, HAA1, ADE8, ADE17 and DOT1) from the prototrophic collection to compare our co-culture screening result with their behavior when grown in bacterial supernatant media. *Pseudomonas* supernatants were prepared in 500 mL of SALN media where the *Pseudomonas sSpp*. strains were inoculated at 0.05 optical density (OD) and incubated for two days at 22°C without agitation. The supernatant of *S. cerevisiae hoΔ::KanMX* was included as a control. Supernatant were filtered with a 0.02 µM membrane and stored at -20°C before use. Each deletion strain was individually grown overnight at 30°C without agitation in SALN. Then, they were inoculated in the supernatants for the growth experiment. Initial OD was adjusted for each strain at 0.05 in 200 µL in a 96 well microplate. Each condition was performed in triplicates. OD was measured each 30 minutes for 60 hours at 30°C using a Tecan plate reader (Zürich, Switzerland). After modelling the growth curve for each well using a Gompertz model fit in JMP13 software, the intrinsic fitness value was calculated by integrating the area under the curve up to 70 hours of growth.

## Data availability

Raw sequencing data are available at Bioproject number PRJNA634859 at http://www.ncbi.nlm.nih.gov/bioproject/. *Pseudomonas* strain 16S rRNA gene sequences are available under the accession numbers MT536139 to MT536145 and RPO-B gene under (numbers to be provided).

## Acknowledgements

We are grateful to the maple syrup producers who provided samples for strain isolation.

## Contributions

Study conception and design: MF, CRL, GN

Acquisition of data: NG, MJ

Analysis and interpretation of data: MF, GN

Drafting of manuscript: MF, GN

Critical revision: MF, GN, CRL, MJ

## Footnotes

Supplementary Information accompanies this paper on Journal website

## Supplementary File Description

**Supplementary File 1**.

Fitness obtained for each yeast deletion strain in each replicate experiment of the co-culture functional genomic screen.

**Supplementary File 2**.

Compilation of enrichment analysis using the online tool of the String Protein-Protein Interaction Networks database.

**Supplementary File 3**.

List of indexes used for sequencing library construction and reference barcodes used for the analysis.

